## Supplemental Figures for "Reactive oxygen species prevent lysosome coalescence during PIKfyve inhibition"

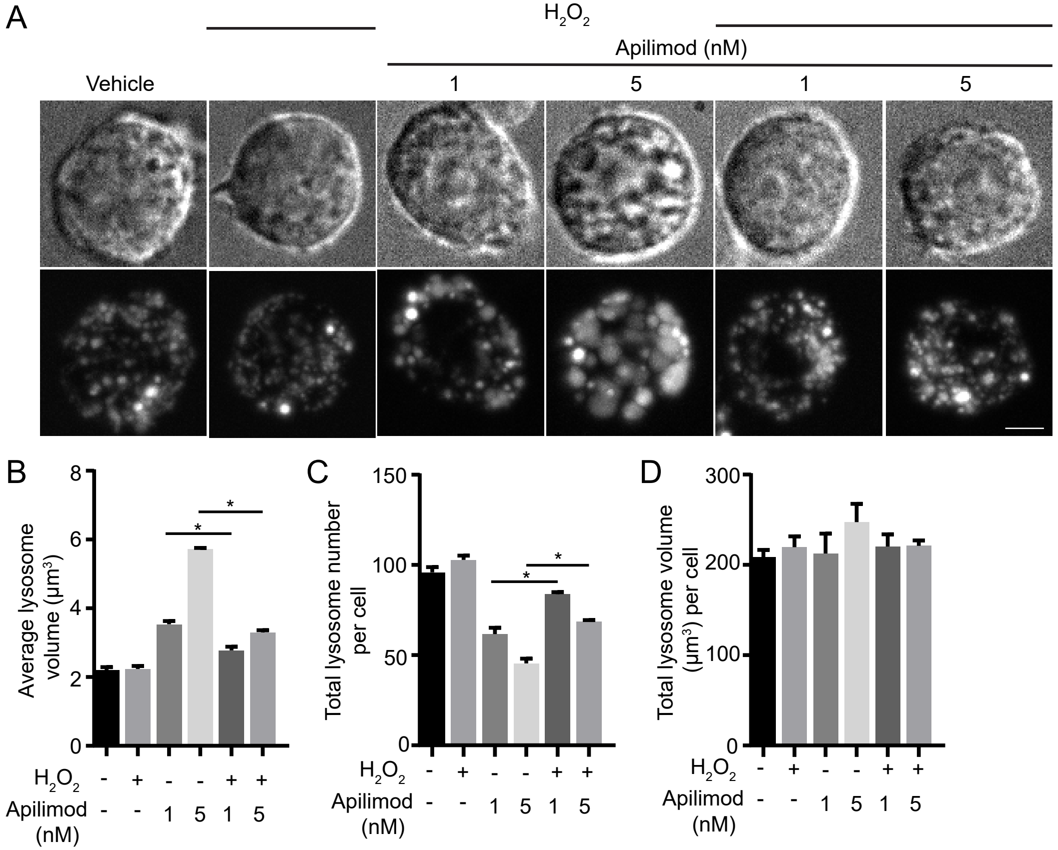


S1 Supplemental Figure

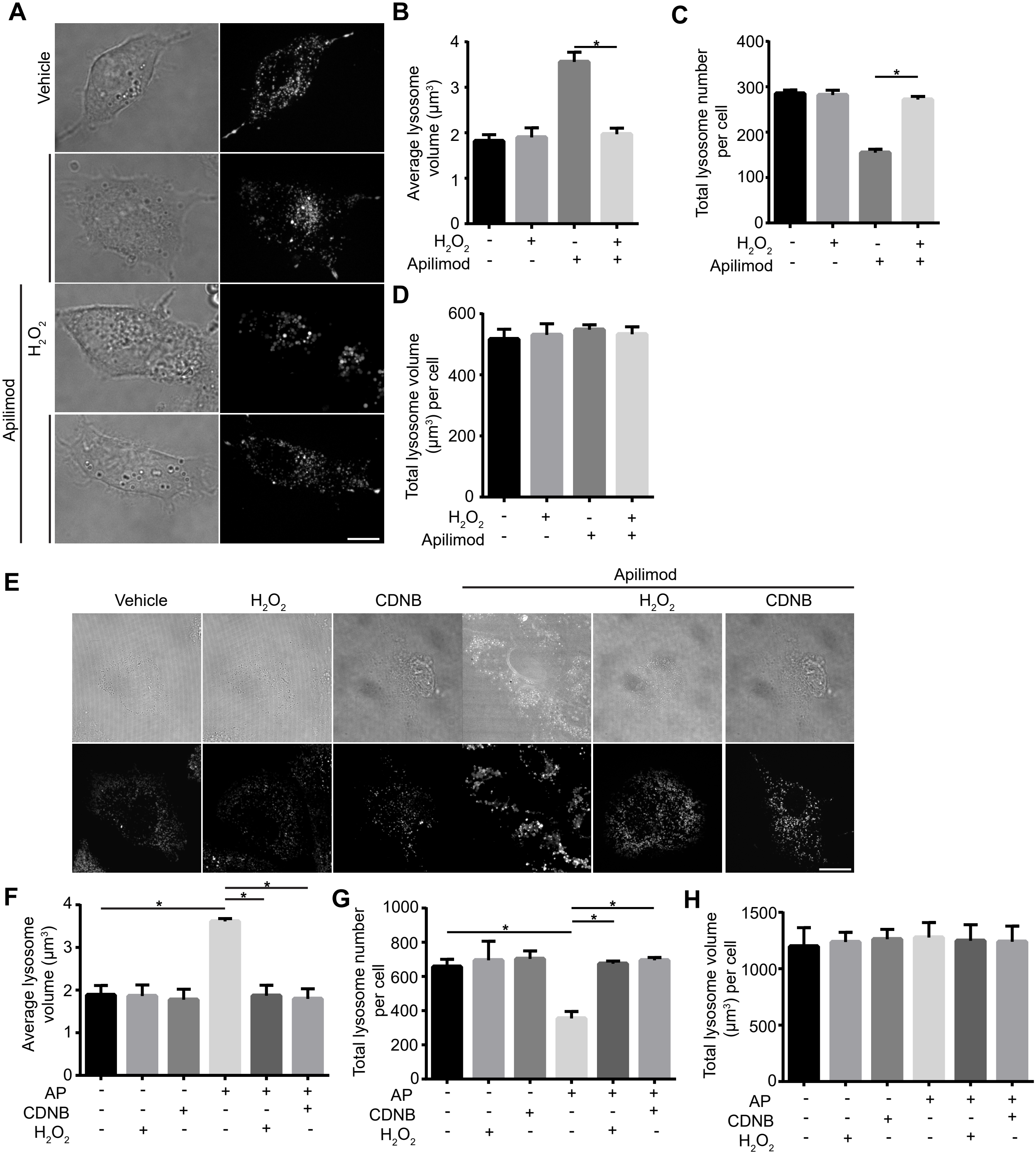


S2 Supplemental Figure

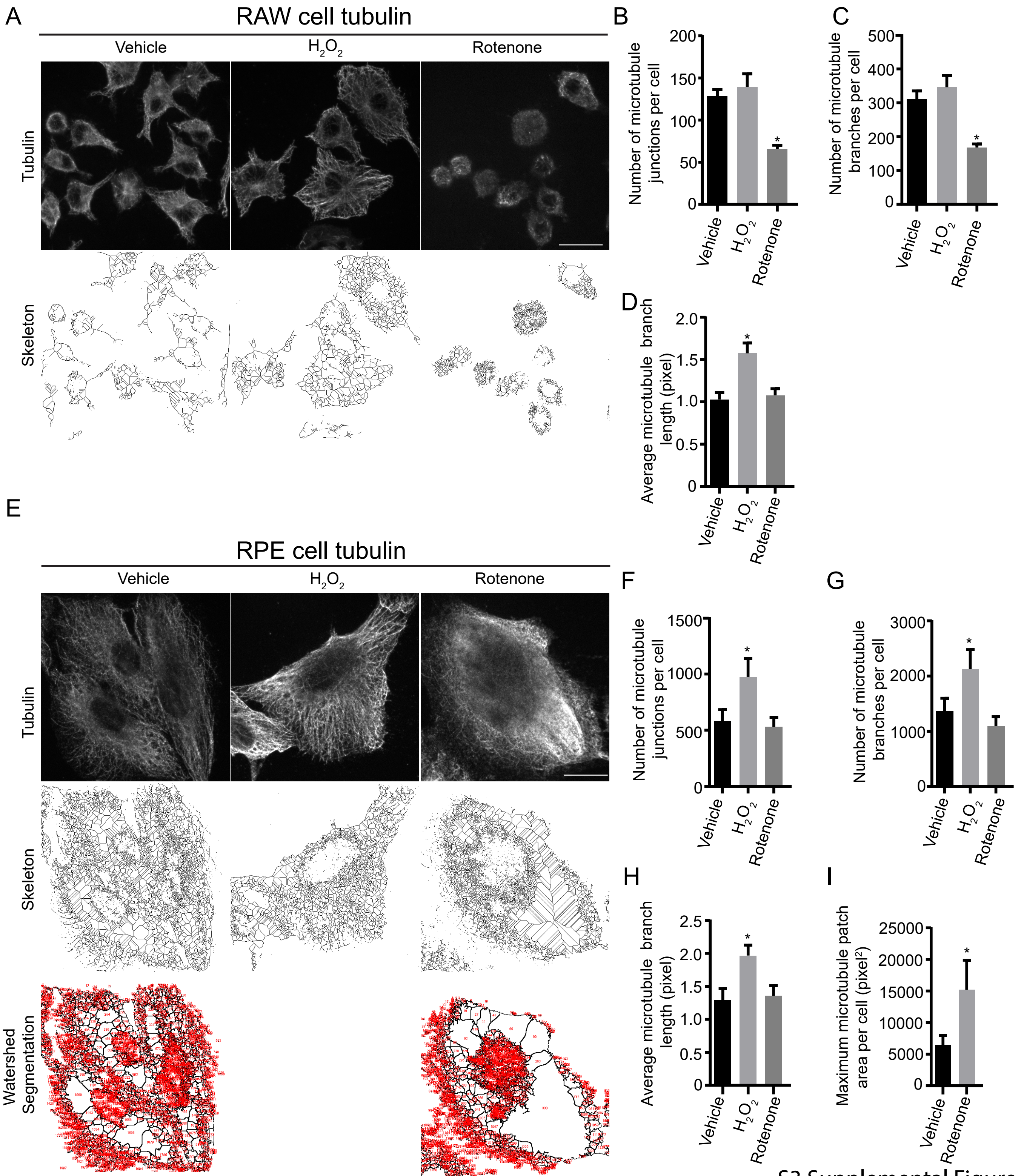


S3 Supplemental Figure

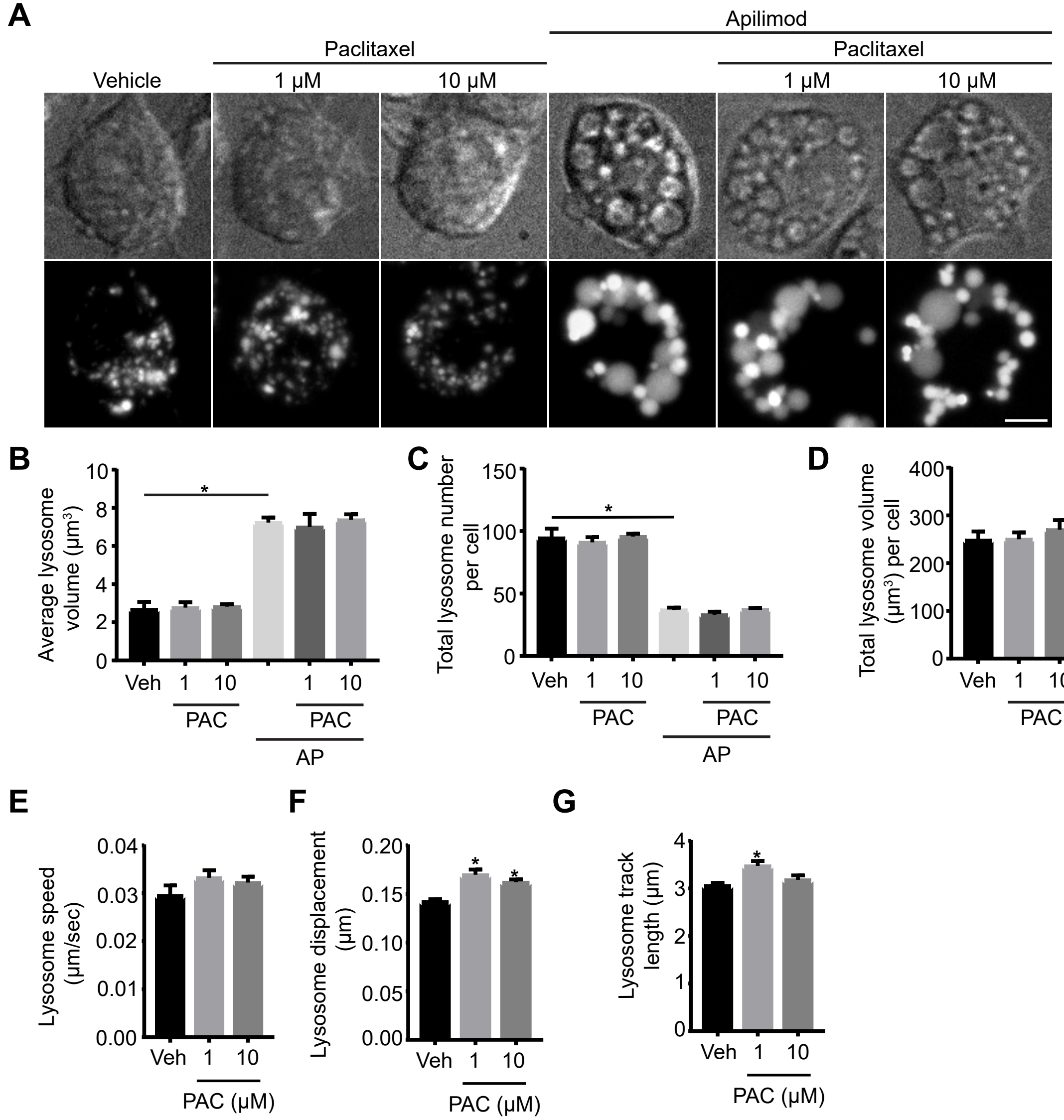


S4 Supplemental Figure

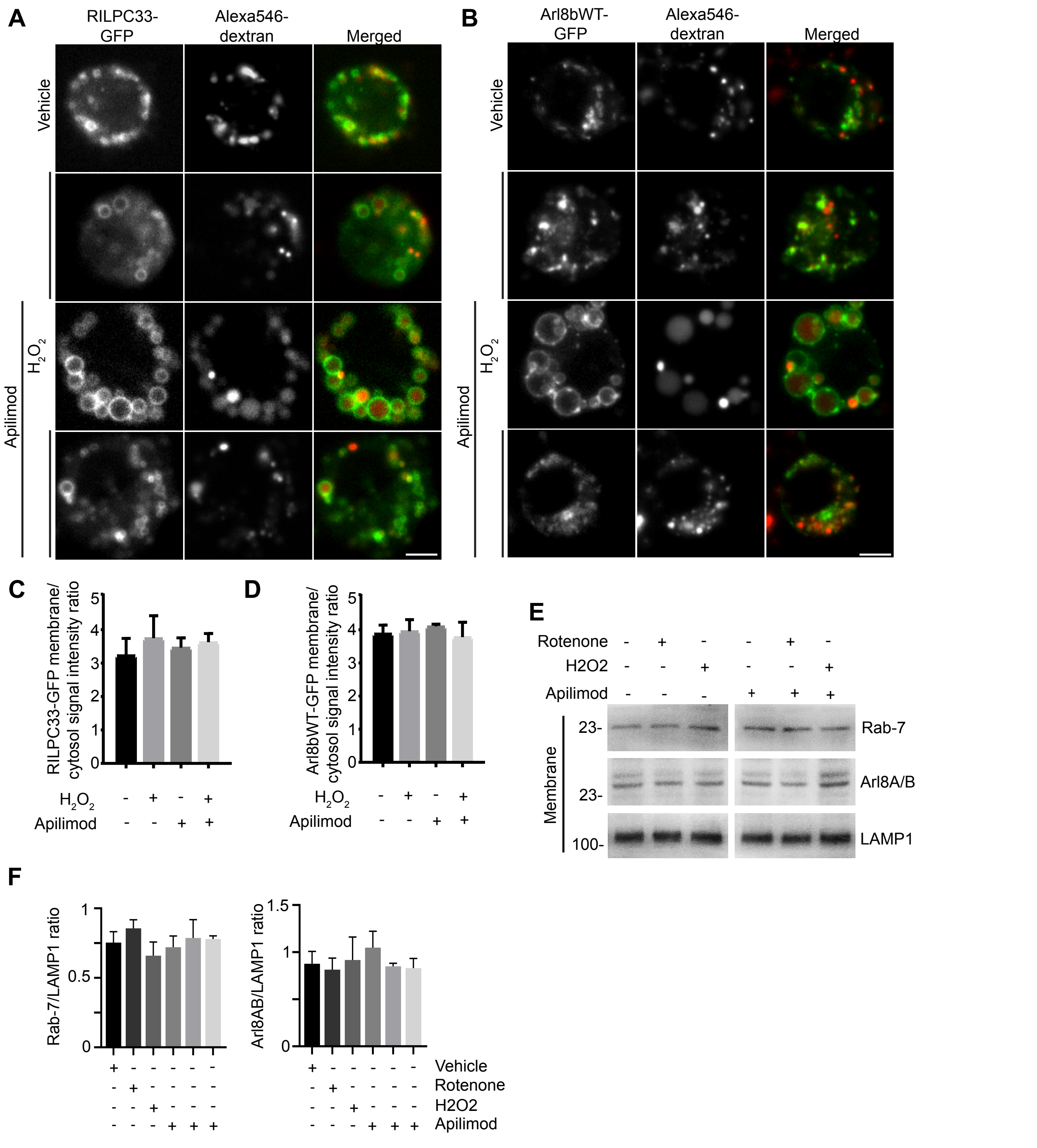


S5 Supplemental Figure

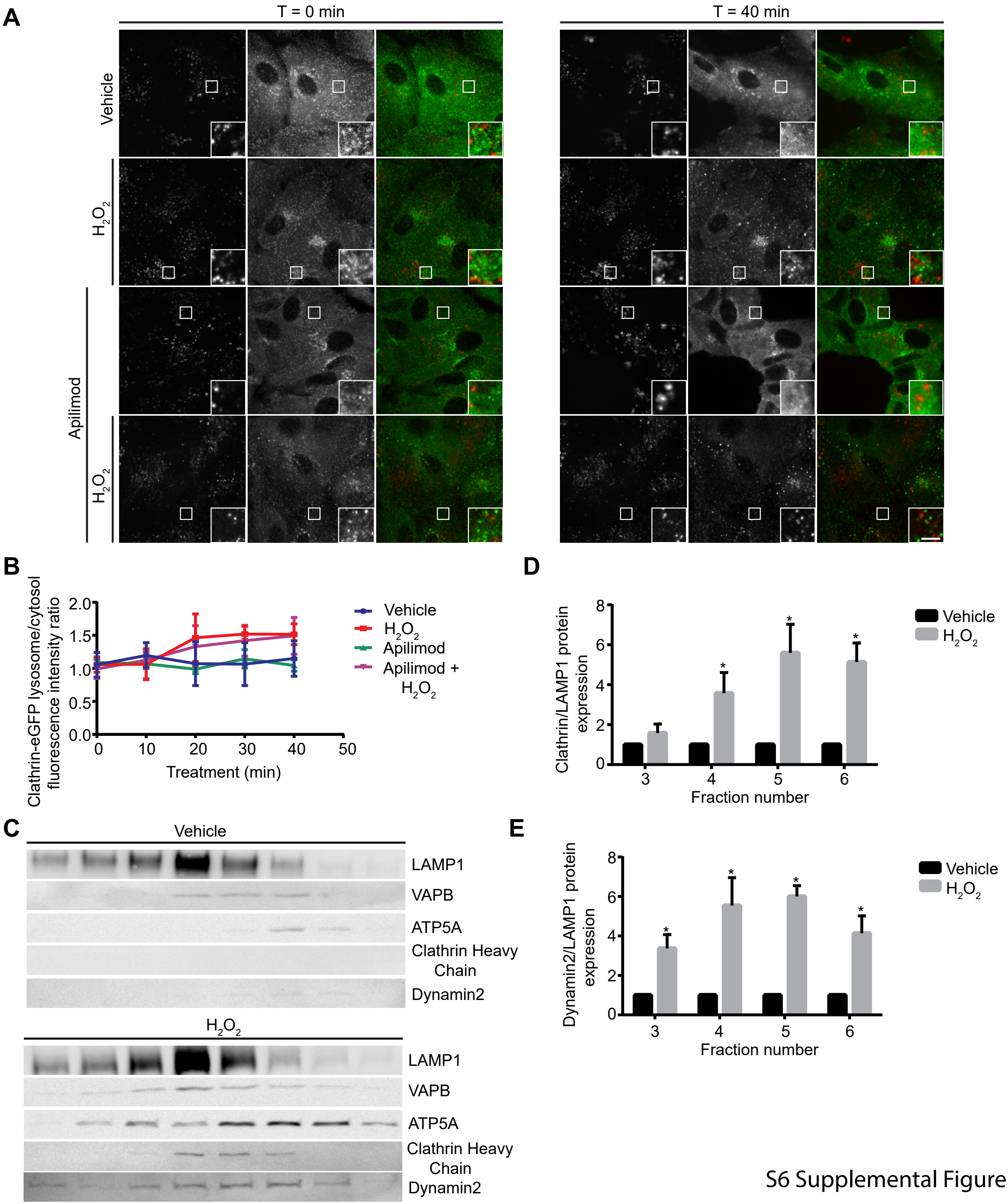


S6 Supplemental Figure

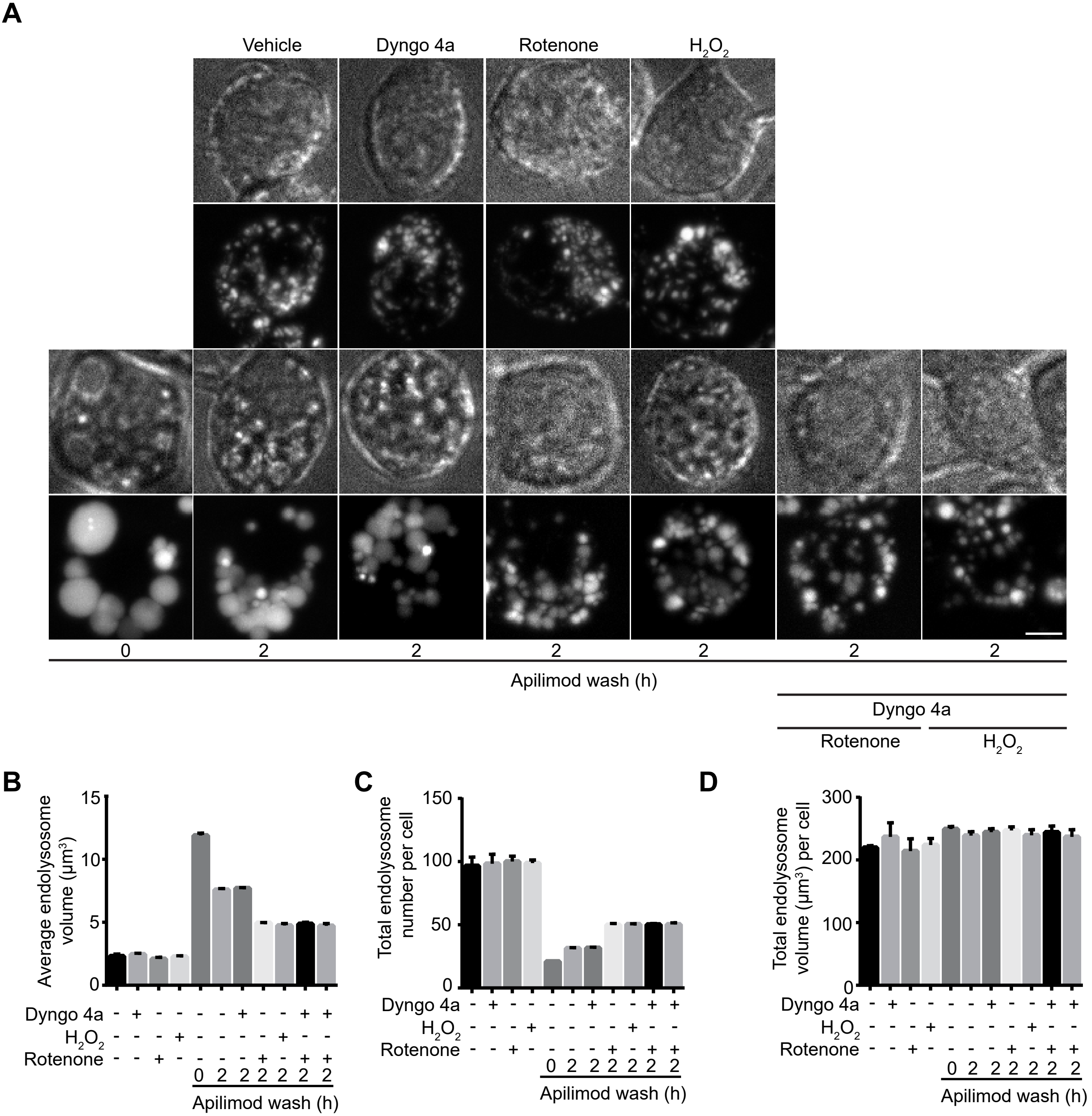


S7 Supplemental Figure
